# Image processing and genome-wide association studies in sunflower identify loci associated with seed-coat characteristics

**DOI:** 10.1101/2021.04.15.439933

**Authors:** Hod Hasson, Mangesh Y. Dudhe, Tali Mandel, Emily Warschefsky, Loren H. Rieseberg, Sariel Hübner

## Abstract

Sunflower seeds (technically achenes) are characterized by a wide spectrum of sizes, shapes, and colors. These traits are genetically correlated with the branching plant architecture loci, which were introgressed into restorer lines to facilitate efficient hybrid production. To break this genetic correlation between branching and seed traits, high resolution mapping of the genes that regulate seed traits is necessary. Recent progress in genomics permits acquisition of comprehensive genotyping data for a large diversity panel, yet a major constraint for exploring the genetic basis of important phenotypes across large diversity panels is the ability to screen and characterize them efficiently. Here, we implement a cost-effective image analysis pipeline to phenotype seed characteristics in a large sunflower diversity panel comprised of 287 individuals that represents most of the genetic variation in cultivated sunflower. A genome-wide association analysis was performed for seed-coat size and shape traits and significant signals were identified around genes regulating phytohormone activity. In addition, significant seed-coat color QTLs were identified and candidate genes that effect pigmentation were detected including a phytomelanin regulating gene on chromosome 17. Finally, QTLs associated with the seed-coat striped pattern were identified and phytohormone regulating candidate genes were detected. The implementation of image analysis phenotyping for GWAS allowed efficient screening of a large diversity panel and identification of valuable genetic factors effecting seed characteristics at the finest resolution to date.

## Introduction

Sunflower (*Helianthus annuus* L.) was domesticated approximately 4,000 years ago in North America (Harter et al. 2004). Since its domestication, selection and breeding has transformed the highly branched, multi-headed, self-incompatible sunflowers into an unbranched, self-compatible crop, with large seeds (technically achenes) that are packaged into a single flowering head. Following domestication by Native Americans, domesticated sunflowers experienced a complex history of breeding which included both divergence from its wild ancestor and subsequent introgressions with wild species to re-introduce valued traits (Putt et al. 1997; Baute et al. 2015). Nowadays, sunflower is cultivated across more than 25 million hectares around the globe (FAO, 2018), mostly as an oilseed production crop, although cultivation for other purposes including confectionery consumption, bird and pet feed, and ornamental uses is also common (Hladni et al. 2017).

Wild sunflowers display a wide spectrum of seed shape and color which are associated with the level of habitat disturbance and particularly the soil characteristics (Ostevik et al. 2016), and several QTLs underlying seed traits were identified (Huang et al. 2020; Todesco et al. 2020). Domesticated sunflower seeds are also characterized by a wide range of sizes, geometric shapes, weight, color pattern, and so forth, and QTLs underlying this variation have been mapped to several different linkage groups in biparental populations and diversity panel (Tang et al. 2006, Reinert et al. 2020). Many of the QTLs were identified next to genomic regions that are associated with domestication syndrome related traits (Burke et al. 2002; Tang et al. 2006; Yue et al. 2009; Reinert et al. 2020). For example, seed characteristics are tightly linked to QTLs regulating the branching phenotype (Tang et al. 2006; Bachlava et al. 2010; Reinert et al. 2020) which was introgressed into the cultivated gene pool from wild sunflower to increase pollen availability during hybrid variety production (Miller and Fick 1997). Although, seed traits were targeted by selection over many generations in accordance with purpose and use, the genetic correlation with plant architecture and other linked or pleiotropic traits remained. Seed size and thickness are correlated with seed weight, oil concentration and yield; thus, identification of candidate genes that regulate these traits has high economic and breeding value. Another defining trait in sunflower is seed-coat color which can vary from black to white through gray or brown and include a specific stripe pattern that can be used as a marker for varieties identification (Aliiev 2020). So far, efforts to map seed-associated QTLs have yielded a relatively low mapping resolution and, in most cases, did not permit identification of candidate genes that regulate the size, shape and color features of the sunflower seed-coat (Tang et al. 2006).

Advances in sequencing and genotyping technologies enable screening of large diversity panels for genetic variation at unprecedented depth and resolution. Along with increases in computing power, genome scale analysis is now feasible for very large datasets, which significantly enhance the power of statistical models to identify the genetic basis of complex traits. Among the most popular approaches to study the genetic basis of quantitative traits are genome-wide association studies (GWAS), which are particularly suitable for high resolution genotype data of large diversity panels characterized by a fast decay in linkage disequilibrium. Despite their many benefits, GWAS have some pitfalls that can increase the rate of false positive associations between genotypes and phenotypes. For example, population structure and kinship among individuals in the diversity panel can lead to spurious associations, especially if the traits of interest are correlated with the history of the represented pedigrees (Korte and Ashley 2013). Another major caveat of GWAS is the inflation in significant associations due to the multiple testing nature of the model; thus, correction for *p*-values is necessary, although in many cases it leads to erosion in true association signals.

Despite the quantum leap in genotyping technologies in the past two decades, phenotyping remained a laborious and limited data collection process. Thus, increasing the number of individuals screened for in depth phenotyping is a limiting factor in gaining statistical power to identify the genes contributing to complex traits of interest at high resolution (Furbank and Tester 2011). Recent investments in automatic phenotyping platforms allow screening of plants continuously with high precision, yet conducting large experiments with thousands of individuals remains a costly and laborious procedure. Image analysis platforms are a promising direction for active research because ample phenotypic data can be generated efficiently by processing pictures of many individuals in a short amount of time (e.g. (Klukas et al. 2014; Knecht et al. 2016)).

Here, we report on the analysis of seed traits in a diversity panel comprised of 287 accessions, which represent circa 90% of the genetic variation in cultivated sunflower (Mandel et al. 2011, 2013). This diversity panel includes inbred lines, historically important open-pollinated varieties (OPVs), and landraces, that could be assigned to four distinct breeding groups: maintainer or restorer oil lines (HA-Oil or RHA-Oil), and maintainer or restorer non-oil lines (HA-nonOil or RHA-nonOil). The entire collection was previously genotyped by whole genome shotgun sequencing and was used in GWAS (Hübner et al. 2019; Gao et al. 2019; Reinert et al. 2020; Todesco et al. 2020); thus this panel is an ideal platform to study the genetic basis of seed-coat characteristics in cultivated sunflower. Combining an image processing phenotyping platform and genome-wide association mapping approach allowed us to identify candidate genes that regulate seed characteristics including phytohormone and pigment regulation.

## Materials and Methods

### Phenotyping seed traits in the SAM population

Seeds of the 287 lines that comprise the cultivated sunflower association mapping (SAM) population were obtained from the USDA National Plant Germplasm System. One seed from each genotype was grown under long day regime (16/8 hours light/dark) and constant temperature of 25°C in a 5-liter pot containing commercial soil mix (“GREEN 90”, Ben-Ari Ltd.). Seeds were collected from each genotype and were used in the experiment. Six seeds of each genotype, corresponding to six replicates, were selected randomly for phenotyping using a flatbed scanner (Epson Perfection V800). Color images of each seed from each accession were recorded in JPEG format at 48-bit with 1200 resolution. The continuous auto-exposure setting and all other performance enhancement properties of the scanner program were disabled. The top of the scanner was disassembled and seeds were scanned in a dark room to avoid shading in the images. Accessions with segregating phenotypes, missing or damaged seeds were removed from the analysis; thus, a total of 265 individuals were screened and analyzed. Image analysis was conducted in ImageJ v.1.52a (Schneider et al. 2012) with a global scale resolution of 47.03 pixels/mm, and the “analyze particles” command with a minimum area of 10mm^2^. Seed shape and area descriptors including length (major axis), width (minor axis), aspect ratio (length/width), perimeter, and roundness were obtained for each seed and averaged per accession. Color measurement including RGB (red, green, blue) and brightness were measured using built-in functions in ImageJ. To measure the rate of black and white in the seeds and overcome moderate contrast issues, the images were converted to 8-bit and adjusted using the “MaxEntropy white” method. Following the conversion and adjustment, the black and white pixels were counted, and the product of their proportions (P_White_ × P_Black_) was used as an index for stripes ratio. The procedure was set as a macro script in ImageJ so that the process could be repeated for all images automatically.

### Genotype Data

Genotype data for all accessions was recently generated for a large collection of wild and cultivated sunflower (Todesco et al. 2020). As part of this project, the entire SAM population was genotyped using the GATK best practice pipeline and the XRQ v1.0 as a reference genome (Badouin et al. 2017). To avoid false assignment of the SNPs coordinates on the reference genome due to potential mis-ordering of contigs, we validated our results using a remapped genotyping dataset to a newer version of the XRQ reference genome (Todesco et al. 2020). The variant calls dataset was further filtered to remove indels, minimum minor allele frequency of 5% and maximum 10% missing data. Next, the filtered SNP dataset was phased and imputed using the software Beagle v.5.0. (Browning et al. 2018) with default parameters and a minor allele frequency filter of 5% was applied again. Linkage disequilibrium decay in the SAM population and for each breeding group was estimated using PopLDdecay v.3.3 (Zhang et al. 2018). To correct for population stratification, a PCA was conducted using an LD-pruned dataset at LD = 0.2 in the R package SNPRealte (Zheng et al. 2012).

### Genome-wide association analyses

To test for associations between genotypes and phenotypes we used three different models as implemented in the software GEMMA v.0.98 (Zhou and Stephens 2012). GEMMA is an efficient platform which allows analyses of millions of SNPs called over hundreds of accessions in a reasonable amount of time on a modest computing platform. Seed phenotypes were included in the models without adjustments except in the multivariate models where GEMMA construct a composite vector from a list of traits.

The models were corrected for population structure using the calculated PCs as covariates, and the genetic relatedness among genotypes was estimated as a centered relatedness matrix in GEMMA. Each measured or calculated seed trait was analyzed independently using a univariate linear mixed model (LMM). To further investigate the association between genotypes and composite seed traits, a multivariate linear mixed model (mvLMM) was conducted for different combinations of measurements that together improve the characterization of the seed color or shape traits. Statistical significance for each SNP was determined using the Wald test and *p*-values were corrected for multiple testing with the SimpleM algorithm (Gao 2011) and the Bonferroni correction. For each model a quantile-quantile (QQ) plot was generated using the package qqman (Turner D. Stephan 2018). Finally, a Bayesian sparse linear mixed model (BSLMM) was conducted to identify polygenic major sparse effect in each trait. This approach is a hybridization between the LMM and the sparse regression and allows more freedom in modeling the genetic architecture that best predicts the phenotypes while controlling for population structure (Zhou et al. 2013).

SNP heritability (also called ‘chip heritability’) was computed for each trait association analysis separately to avoid a bias in estimating the percent of the phenotypic variance explained by SNPs detected from a combination of traits. Thus, the percent of variance explained (PVE) was calculated using both the LMM and BSLMM models.

In addition, the percent of genetic variation explained (PGE) by the sparse effect was estimated in the BSLMM model to account for the contribution of the detected significantly contributing SNPs from the overall genetic variation observed for each trait. The calculation of the PVE estimates differ between the LMM and BSLMM, yet the latter is considered more robust to a wider spectrum of genetic scenarios (Zhou et al. 2013).

To identify candidate genes within significantly associated regions, LD was calculated around the top SNP at each region and for each trait. The intervals at both sides of the SNP were searched using the XRQ genome browser (https://www.heliagene.org/HanXRQ-SUNRISE/) until LD dropped below *r^2^* = 0.5.

## Results

### Seed coat characterization in the sunflower association mapping population

A wide range of phenotypic variation was observed across the SAM population for seed-coat characteristics. The seed-coat area ranged from 15mm^2^ to 119mm^2^ and was significantly higher (*w* = 7384, *p* = 9.09×10^-7^) in maintainer lines (n=109) compared with restorer lines (n = 97), and in non-oil types (n = 69) compared with oil types (n=137) (*w* = 3158, *p* = 1.04×10^-4^). The seed coat shape, measured by aspect ratio, roundness (0-1, 1 = perfect circle) and the seed coat perimeter (mm) were also highly variable among accessions with significant differences between breeding groups (Figure 1, Table 1). Overall, maintainer lines are characterized by larger and rounder seed-coats than restorer lines, and non-oil varieties display larger seed-coats than oil type varieties (Figure 1, Table S1). These results for seed-coat size and shape illustrate the differences between breeding material in accordance with the plant architecture and use; branched restorer lines with multiple flowers have smaller seeds, and larger seeds are usually favored for confection consumption.

**Figure 1.**
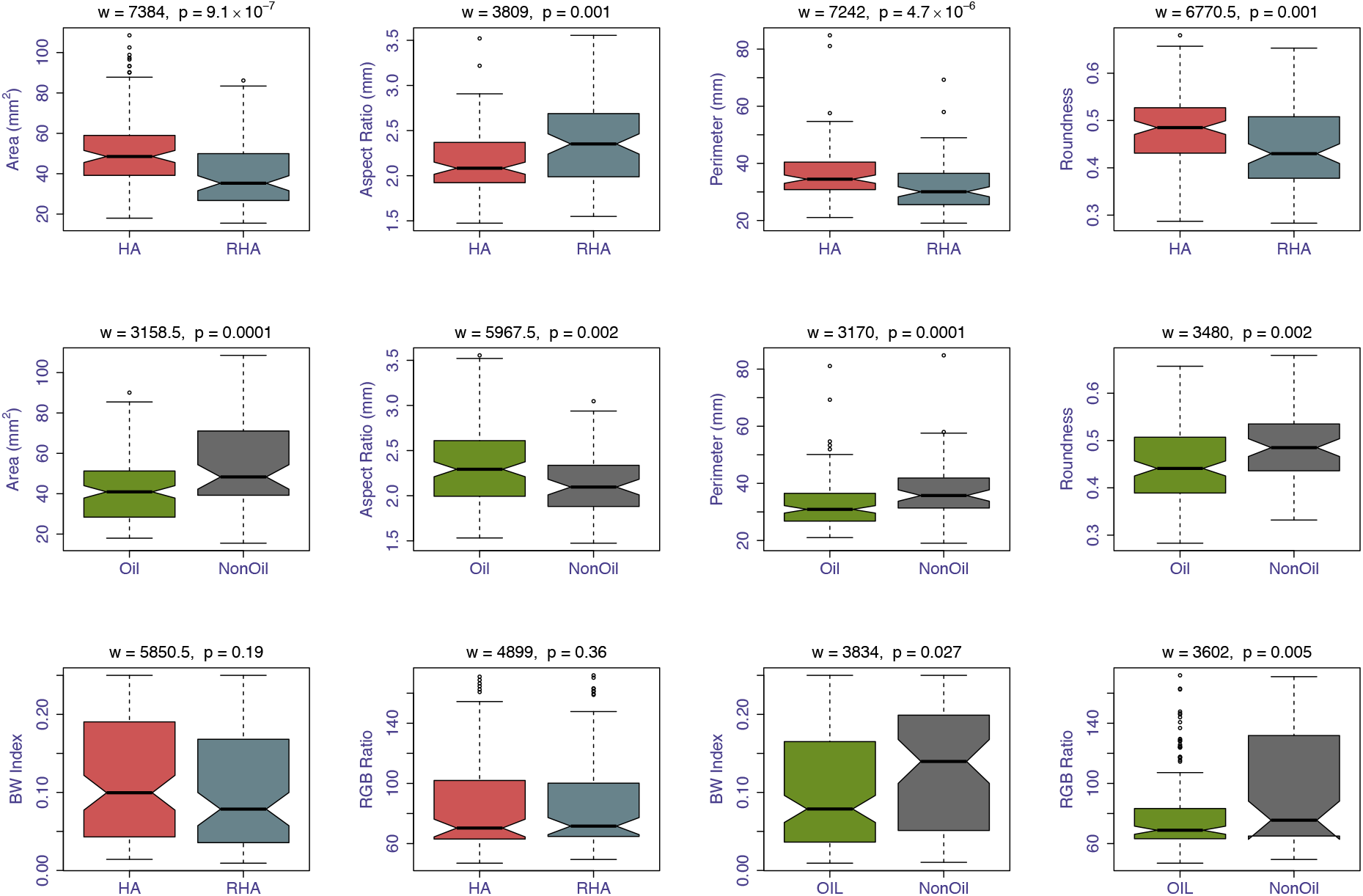
Seed-coat size and color features in maintainer lines (blue, HA; n=109) compared with restorer lines (red, RHA; n = 97), and in oil types (green; n = 137) compared with non-oil types (grey; n=69). Statistical analysis was conducted with the non-parametric Wilcoxon test and *p*-values are indicated at the top of each plot. At the top two rows are the results for seed-coat size and shape characteristics and at the bottom row are color (RGB) and stripe pattern (BW index).

**Table 1.** Summary table of seed-coat characteristics and heritability. Listed are the seed-coat area, aspect ratio, brightness, red-green-blue (RGB) ratio, black-white (BW) index, major and minor axes and the seed-coat perimeter. For each trait, the mean, median, range and the percent of explained variance (PVE) estimated from the univariate LMM and Bayesian sparse LMM models are provided. For heritability scores calculated with the BSLMM, the percent of genotype effect (PGE), which corresponds to the proportion of genetic variance explained by the sparse effect terms, and the average number of SNPs with major effect (N), are also provided.

| Trait | Mean (SD) | Median | Range | LMM |  | BSLMM |  |  |
| --- | --- | --- | --- | --- | --- | --- | --- | --- |
|  |  |  |  | PVE | SE | PVE | PGE | N |
| Area | 47.74 (20.39) | 45.79 | 15.49 - 119.09 | 13 | 9 | 39 | 48 | 118 |
| Aspect Ratio | 2.22 (0.42) | 2.15 | 1.229 - 3.555 | 0.00064 | 13 | 24 | 48 | 17 |
| Brightness | 96 (39.34) | 76.46 | 49.74 - 188.88 | 16 | 9 | 31 | 73 | 7.2 |
| RGB ratio | 86.27 (32.53) | 71.38 | 46.95-171.85 | 17 | 9 | 33 | 79 | 9 |
| BW Index | 0.11 (0.08) | 0.088 | 0.0093 - 0.25 | 10 | 9 | 21 | 43 | 103 |
| Major Axis | 11.08 (1.71) | 10.84 | 7.496 - 17.03 | 16 | 9 | 39 | 51 | 14 |
| Minor Axis | 5.3 (1.5) | 5.23 | 2.620 - 9.931 | 2 | 8 | 33 | 32 | 19 |
| Perimeter | 34.53 (9.56) | 32.81 | 19.05 - 84.82 | 3 | 7 | 26 | 65 | 10 |

To explore the phenotypic variation in seed-coat color, the RGB ratio (red, green, blue) and brightness were calculated from each seed image. In addition, the numbers of black or white pixels were counted and their proportion was used as an indication for the color pattern on the seed-coat (Figure 1, Table 1). Most seeds were characterized by a grey-black color (mean_RGB_ratio_ = 86.27, SD_RGB_ratio_ = 32.52) and uniform pattern (mean_BW_index_ = 0.11, SD_BW_index_ = 0.08). The brightness value was significantly correlated with the RGB ratio (*r* = 0.993, *p* = 2.2×10^-16^), indicating these traits are equivalent for sunflower seed-coat characterization (Figure 2). Interestingly, a significant correlation (*r* = 0.44, *p* = 3.57×10^-14^) was observed between the seed-coat color (RGB ratio) and size traits (Figure 2), indicating that larger seeds tend to lose pigmentation and become brighter. It has become clearer when we observed a significant positive correlation (*r* = 0.38, *p* = 1.10×10^-10^) between BW index and seed-coat area; thus, larger seeds are brighter and less uniform (Figure 2).

**Figure 2.**
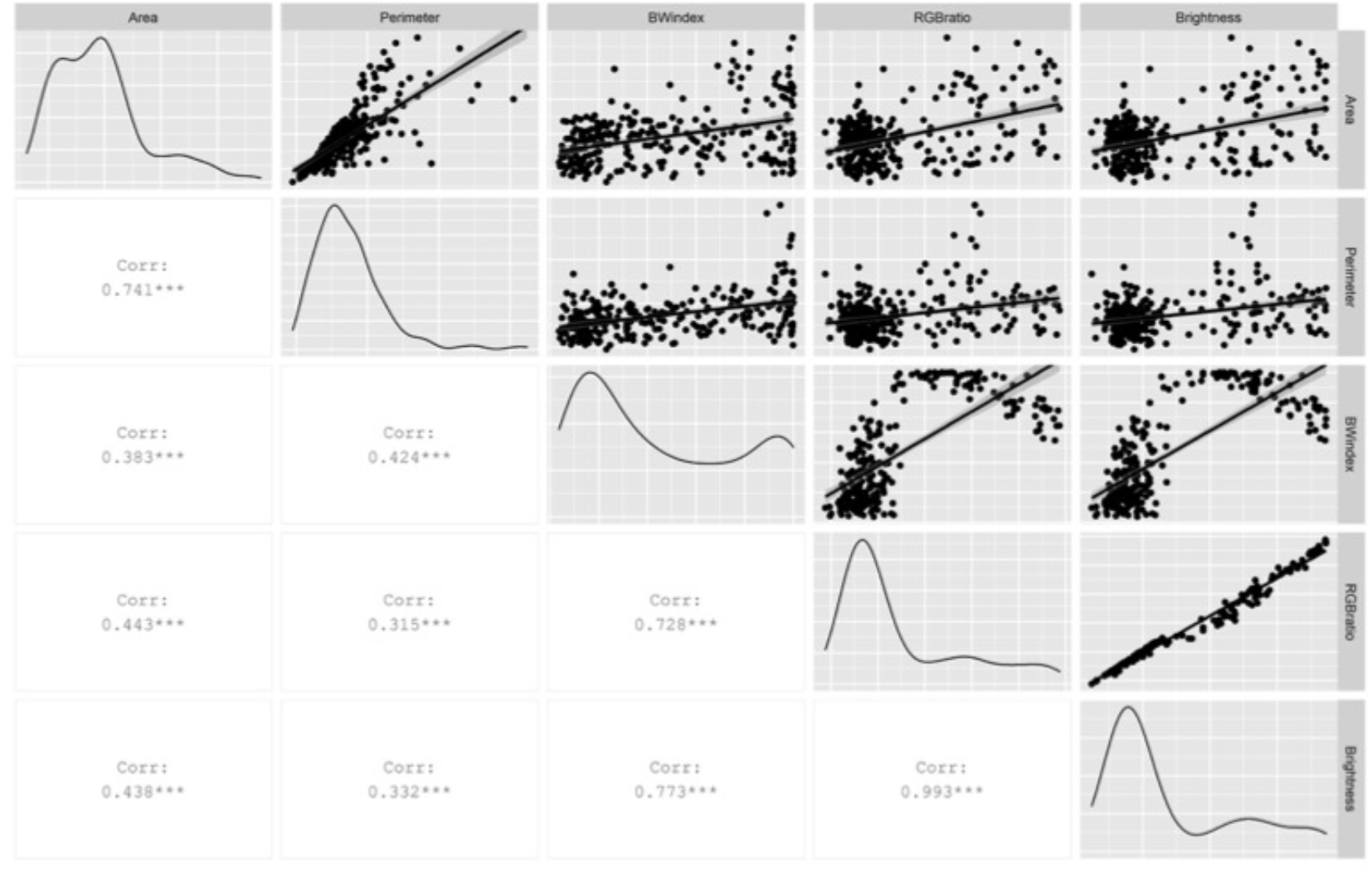
Pearson correlation of seed size and color features. On the diagonal are the traits distribution, on the lower triangle are the correlation coefficients and significance marked with asterisks, and on the upper diagonal are the scatter plots for each pair of traits.

### Exploring the genetic basis of seed-coat characteristics

The sunflower association mapping (SAM) population represents approximately 90% of the genetic variation in cultivated sunflower (Mandel et al. 2011, 2013). This diversity panel can be genetically split into four groups (Figure 3) in accordance with the line type (restorer/maintainer) and use (oil/non-oil). To identify genomic regions and candidate genes associated with seed-coat traits, a genome-wide association analysis was performed using three different models. The decay of LD varied between groups, where maintainer non-oil group (HA-NonOil) is characterized by the lowest LD (*r^2^* = 0.5 at 7 Kbp), and restorer non-oil (RHA-NonOil) group has the highest LD (*r^2^* = 0.5 at 60 Kbp). Across the entire SAM population, LD decay reached *r^2^* = 0.50 at 5 Kbp (Figure 3). To correct the effect of population structure, a PCA was conducted on an LD-pruned and imputed dataset and the first four PCs were used as covariates in all statistical models. Adjustment of the significance threshold was set using the SimpleM (1.5×10^-7^) and Bonferroni (3.2×10^-8^) methods to account for multiple comparisons. To identify genomic regions associated with seed-coat shape and color, three statistical models were applied: a univariate model for each trait separately, a multivariate model which considers different combinations of shape and size characteristics as a single composite trait, and a Bayesian sparse linear model which allows to test for concentrated polygenic effects. All analyses were conducted with both SNP dataset to validate the coordinates of the identified QTLs and candidate genes on the XRQ reference genome.

**Figure 3.**
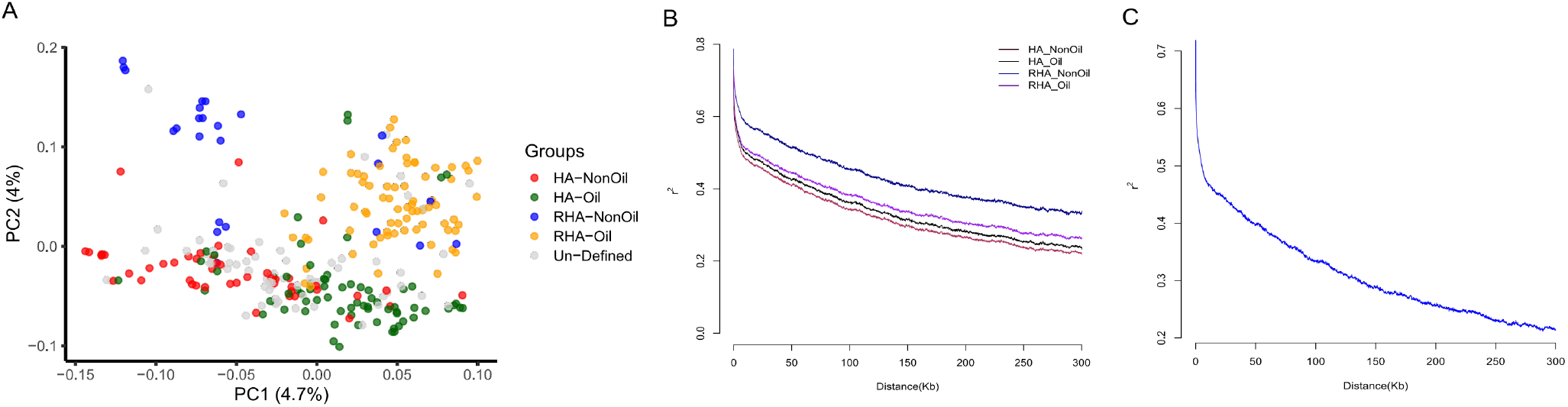
Major sub-populations comprising the sunflower association mapping (SAM) population. A) Principal components analysis for the SAM population based on 1.5 million SNPs. Colors correspond to the known assignment of each accession to a group and unknown assignments are marked in grey. B) Linkage disequilibrium (LD) decay plot conducted for each of the four breeding groups in the SAM population, and C) LD decay across the entire SAM population. Linkage disequilibrium, was calculated in windows of 500Kbp for pairs of polymorphic marker loci.

Significant associations between seed-coat size and shape characteristics were identified by the univariate, multivariate and Bayesian sparse models on chromosome 17 (194,057,300-194,067,000) and chromosome 10 (33,070,000-33,071,000; 187,558,000-188,232,500). The multivariate model for the composite size trait, consisted of the seed-coat area, major and minor axes, and perimeter, and allowed to identify additional significant association signals on chromosomes 13 (118,145,400-118,330,700), and 14 (137,658,300-137,670,400). These results support the significant difference observed in seed size between maintainer (HA) and restorer (RHA) lines, where the latter accommodate the fertility restoration allele on chromosome 13 and are characterized by a branchy plant architecture (Figure 1 and Figure 4).

**Figure 4.**
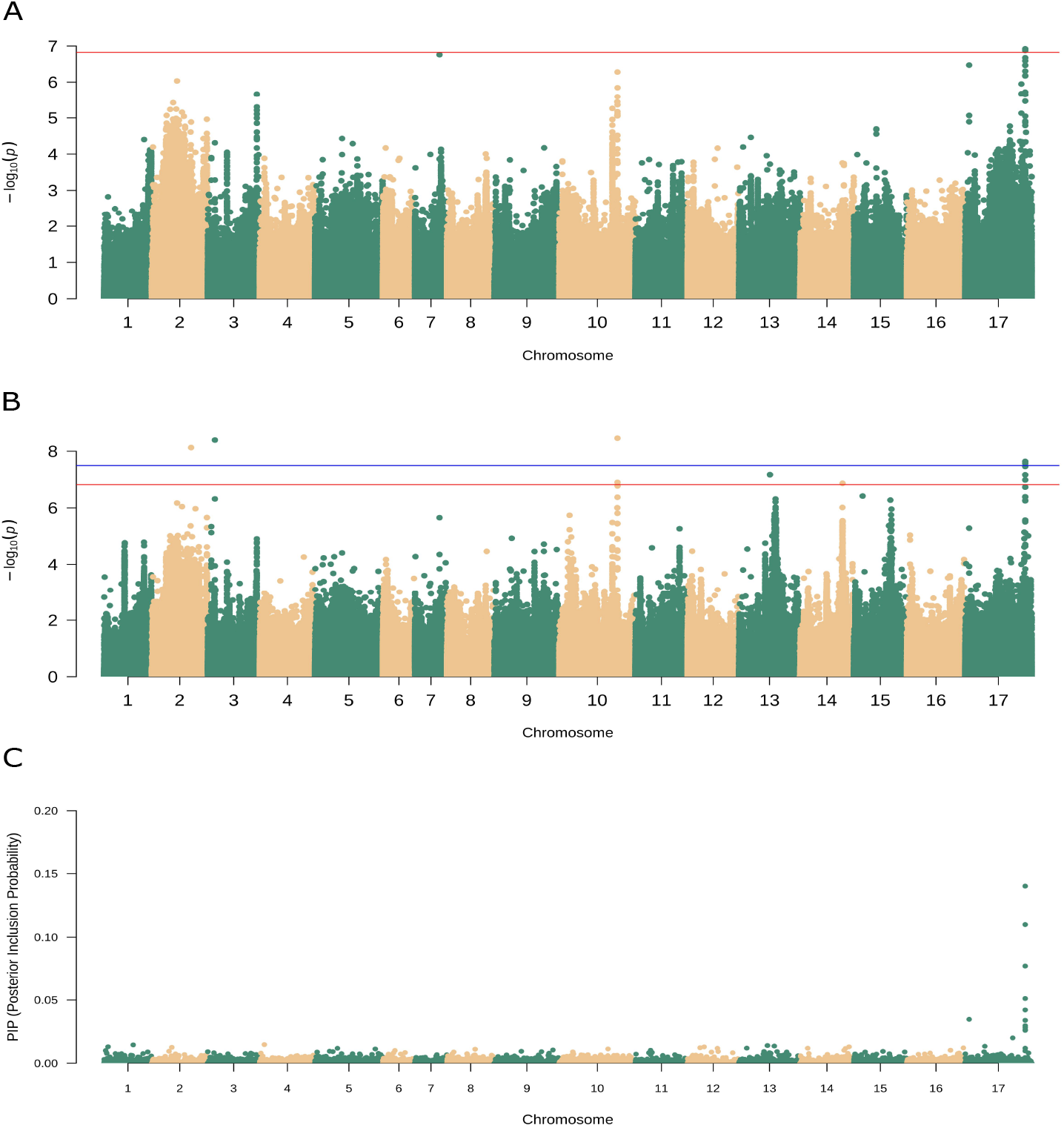
Genome-wide association mapping of seed-coat size and shape traits. On the left are Manhattan plots for the different models where red and blue horizontal lines correspond to the simpleM and Bonferroni correction threshold respectively. On the right of each Manhattan plot are the qq plots for each model. A) Univariate linear model conducted for the major axis trait, B) multivariate linear model analysis which include the seed-coat area, major and minor axes and perimeter, C) a Manhattan plot for the BSLMM model conducted for the major axis trait.

Candidate genes were identified mainly in chromosomes 17 and included genes that regulate cell division and growth through gibberellin regulating genes (HanXRQChr17g0566801) and ethylene regulating genes (HanXRQChr17g0566641, HanXRQChr17g0566651). Other candidate genes that are involved in plant growth include the MOB kinase activator (HanXRQChr17g0537381), which plays a key role in the regulation of cell division and expansion. A complete list of candidate genes and their location is provided in the supplementary information (Table S2).

Association mapping of different color traits based on the univariate and Bayesian sparse models permitted to identify significant and overlapping genomic regions on chromosomes 1 (148,076,600-148,106,300), 14 (142,274,000-142,277,000), and 17 (144,422,500-144,647,000). For the BW index trait, strong associations were observed on chromosomes 11 (10785800-10791500) and 16 (614700-619300), although these signals did not pass the adjusted significance threshold in the univariate model. These association signals were observed also in the multivariate model combining the three color-features: RGB ratio, brightness, and BW index, and to lower extent also in the BSLMM (Figure 5). Candidate genes were identified under the detected regions and included several flavonoid biosynthesis genes on chromosomes 13 and 14 (HanXRQChr14g0448921, HanXRQChr14g0448931, HanXRQChr14g0448941), 16 (HanXRQChr16g0497731) and 17 (HanXRQChr17g0560701), and a major phytomelanin synthesis gene (laccase) on chromosome 17 (HanXRQChr17g0560821). Both pigments were previously described as the main factors effecting the seed-coat color in sunflower (Miller and Fick 1997).

**Figure 5.**
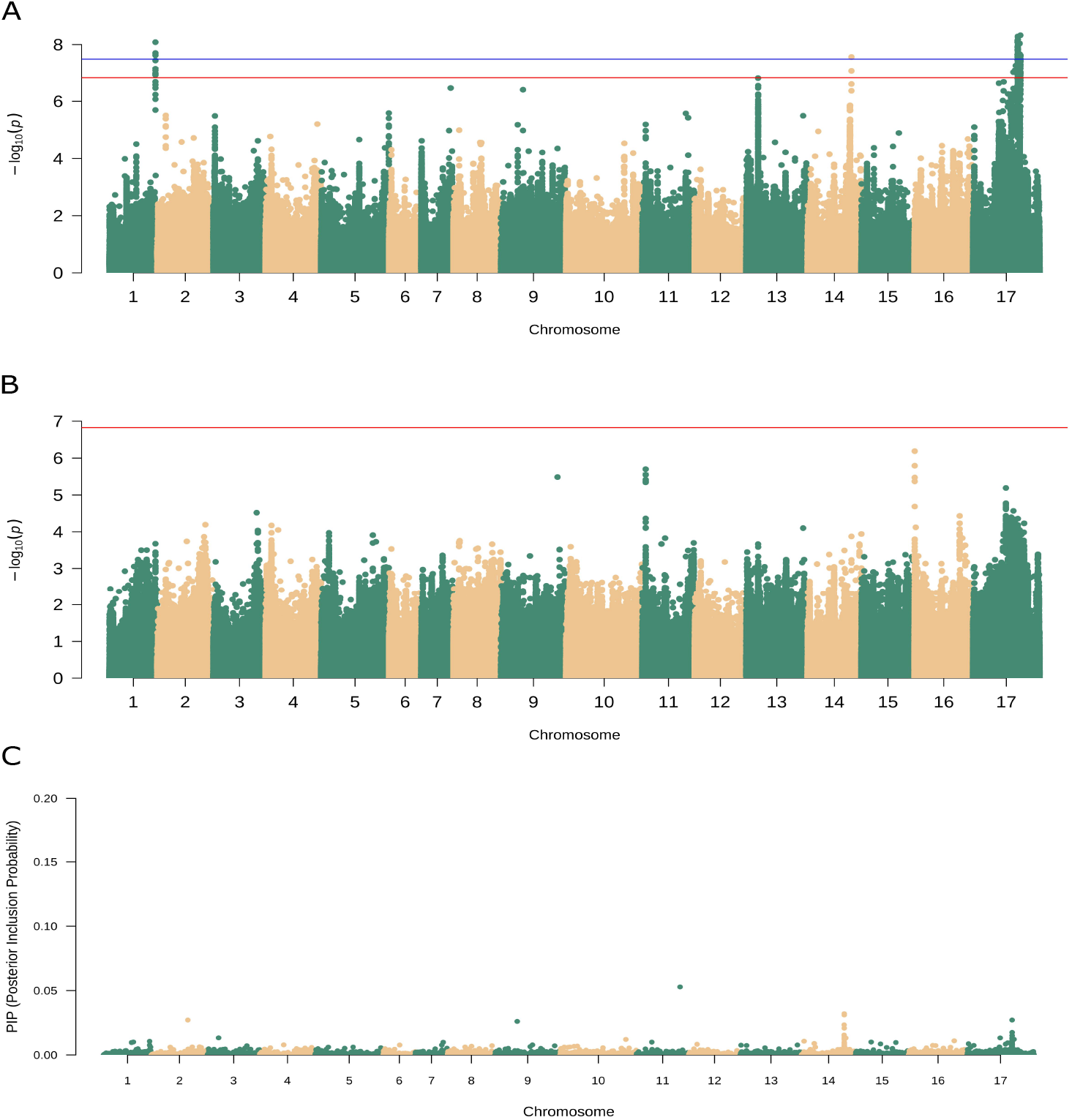
Genome-wide association mapping of seed-coat size and shape traits. On the left are Manhattan plots for the different models where red and blue horizontal lines correspond to the simpleM and Bonferroni correction threshold respectively. On the right of each Manhattan plot presented are qq plots for each model. A) Univariate linear model conducted for the RGB ratio trait, B) univariate linear model conducted for the BW index trait, C) a Manhattan plot for the BSLMM model conducted for the RGB ratio trait.

### The heritability of seed coat traits

Seed coat characteristics have been clearly selected by breeders in accordance with the type and use of the different varieties. While complex quantitative genetic inheritance was recently described for seed size traits (Tang et al. 2006; Reinert et al. 2020), Mendelian inheritance has been proposed for color characteristics and pattern (Gorohivets and Vedmedeva 2016). Nevertheless, little is known about the heritability of these traits and more specifically the SNP heritability (‘chip heritability’), which is used as a proxy for the narrow-sense heritability, a key factor in predicting the response to selection in a breeding program. To test the SNP heritability of seed characteristics, the percent of explained variance was estimated for each trait separately using the results obtained from the univariate model and BSLMM (Table 1). According to BSLMM, the seed-coat major axis and RGB ratio showed the highest heritability values of 39% and 33% respectively, and over 50% of the genetic effect in the major axis trait could be explained by 14 major SNPs that correspond to one region on chromosome 17 (194,055,000-194,070,000) where most of the candidate genes were detected. Similarly, over 75% of the genetic effect in the RGB ratio color trait could be explained by 9 SNPs, of which 4 SNPs were located on chromosome 14 (137,200,000-137,700,000), and two on chromosome 17 (144,450,000-144,500,000).

## Discussion

Sunflower is among the most important crops used for edible oil production, however other uses including confectionery purposes and ornamental use are also common. Seed size, shape and color are considered domestication syndrome traits (Burke et al. 2002), and are associated with the plant branching architecture (Bachlava et al. 2010). Here, we used image processing analysis to characterize seed-coat features and genome-wide association analyses to explore the genetic basis of these key phenotypes.

Overall, seed-coat shape and size were significantly correlated with the variety use; i.e. larger seeds were observed in maintainer lines compared with the branchy restorer lines, and in confectionary varieties compared with oil varieties. These results are in line with previous studies and reflect the importance of these traits in breeding for different purposes (Tang et al. 2006; Bachlava et al. 2010; Hladni et al. 2017). The seed-coat color and pattern were also attributed to purposes of the varieties, where larger and brighter or stripy seed-coats are usually associated with confectionary varieties, and small uniformly colored seed-coats are associated with oil production varieties. The observation that small seeds are darker and large seeds are brighter could potentially reflect the pigmentation density; thus, small seeds are characterized by dense pigmentation compared with large seeds. The Black-White index is indicative of the stripy pattern on the seed-coat, which presumably reflects the coloration in accordance with the fiber orientation along the major axes of the seed-coat.

The SAM population provides an excellent platform to study the genetic basis of quantitative traits using a genome-wide association approach. Overall, seed-coat shape and size overlap known QTLs associated with the maintainer-restorer phenotypes, including the branching locus on chromosomes 10 and fertility restoration locus on chromosome 13 (Bachlava et al. 2010; Mandel et al. 2013; Baute et al. 2015; Hübner et al. 2019). In addition, our analysis revealed a strong signal with high heritability on chromosome 17 by all three models, indicating that seed-coat size is not the outcome of plant architecture exclusively, and that this genomic region is a good target for identification of candidate genes that are specifically involved in seed-coat characteristics. Indeed, several candidate genes that regulate phytohormone biosynthesis were identified on chromosome 17 including genes that regulate cell division and expansion through gibberellin, ethylene and auxin biosynthesis. Previous studies reported a strong correlation between plant architecture and seed size (Burke et al. 2002; Tang et al. 2006), and phytohormone synthesis and regulation are expected to have a major effect on both traits. The high heritability attributed to this region indicates that the associated SNPs could be used as good molecular markers for seed size and presumably also plant architecture breeding.

Genomic regions associated with color traits were observed on chromosomes 1, 11, 14, 16 and 17. A previous study (Gorohivets and Vedmedeva 2016), suggested Mendelian inheritance of two loci that effect the stripy pigmentation pattern on the seed-coat. The GWAS analyses conducted here indeed located two regions on chromosomes 11 and 16. Interestingly, candidate genes found in these regions correspond to gibberellin regulation and auxin-flavanol biosynthesis; thus, it is plausible that phytohormone regulation generates the stripy patterns through differential cell division and elongation. In addition, pigmentation related genes were identified including a phytomelanin regulating gene on chromosome 17.

The high-throughput platform used in this study to phenotype seed-coat characteristics based on image analysis and GWAS represents a promising approach for studying the genetic basis of important traits. This approach is highly efficient and allowed us to identify QTLs associated with traits of interest and the underlying candidate genes. Despite the efficiency of the phenotyping method, more complex traits should be validated under field conditions to guarantee the concordance between image-based analyses and phenotyping for commercial breeding purposes.

## Acknowledgments

We thank Eidit Shtern, Nadav Uzan and Shay Uzan for assistance with phenotyping and the Hübner lab members for discussion and comments on earlier versions of the manuscript.

## Funding

This Research was supported by the Global Crop Diversity Trust grant no: GS17008 and the Israel Ministry of Agriculture grant no: 21-01-0034.

## Notes

### Competing Interest Statement

The authors have declared no competing interest.

